## Supplementary material for "Atlas of *Plasmodium falciparum* intraerythrocytic development using expansion microscopy": Figure Supplements

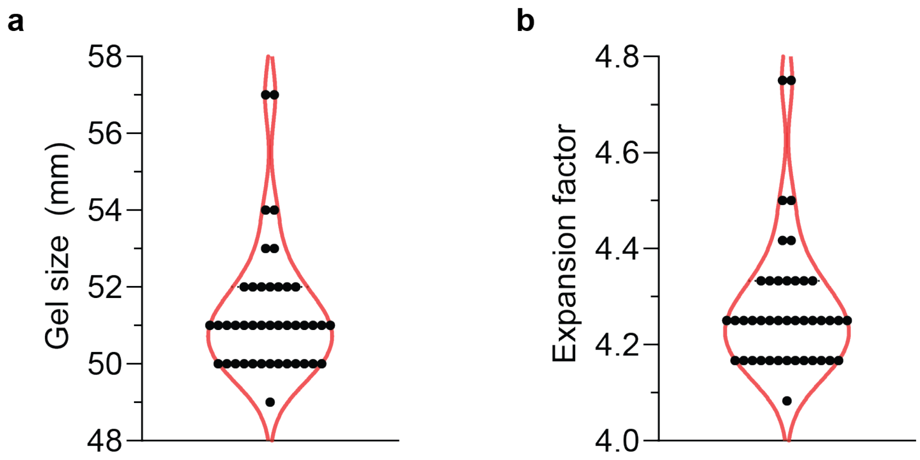


**Figure 1 – Figure Supplement 1**

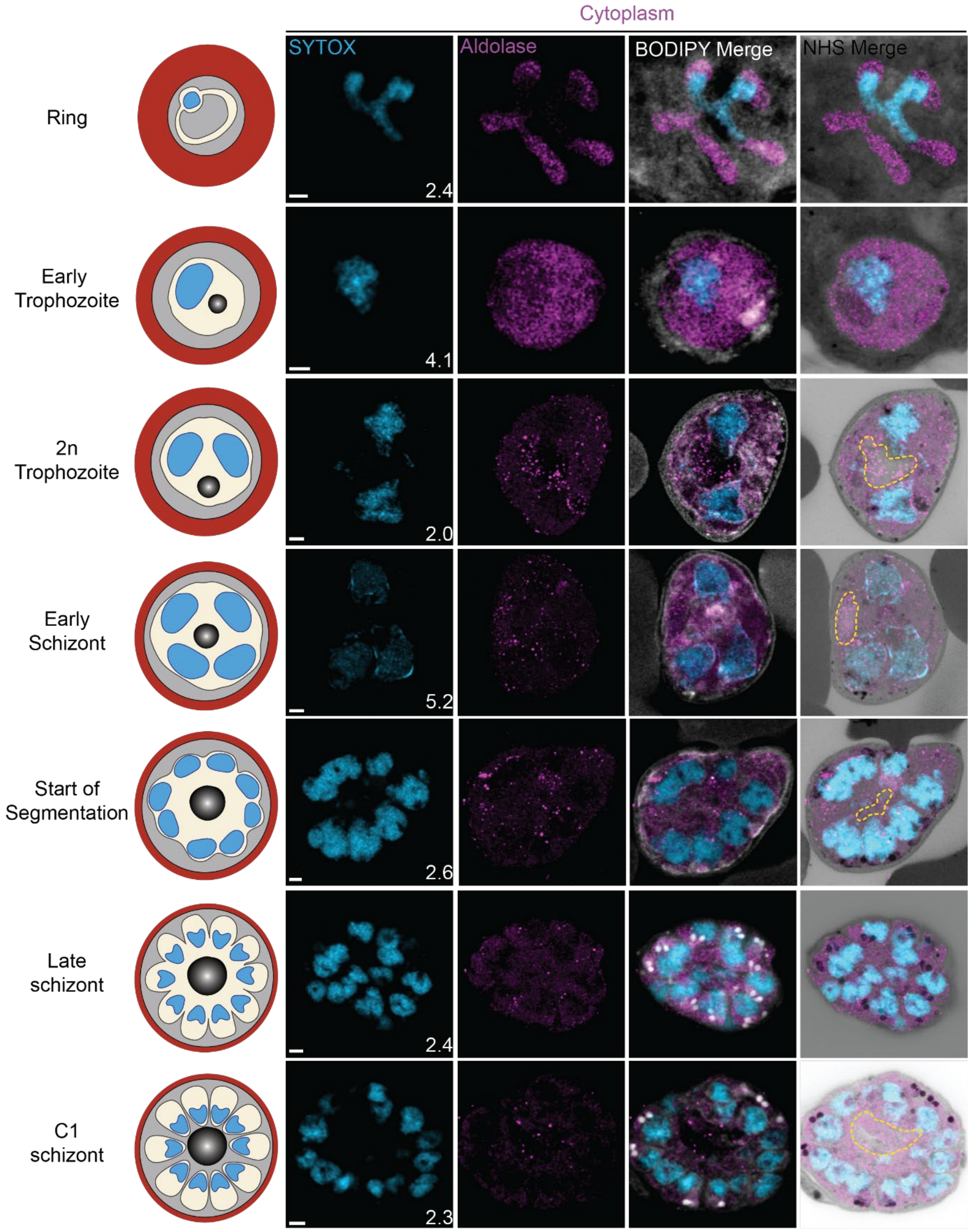


**Figure 1 – Figure Supplement 2**

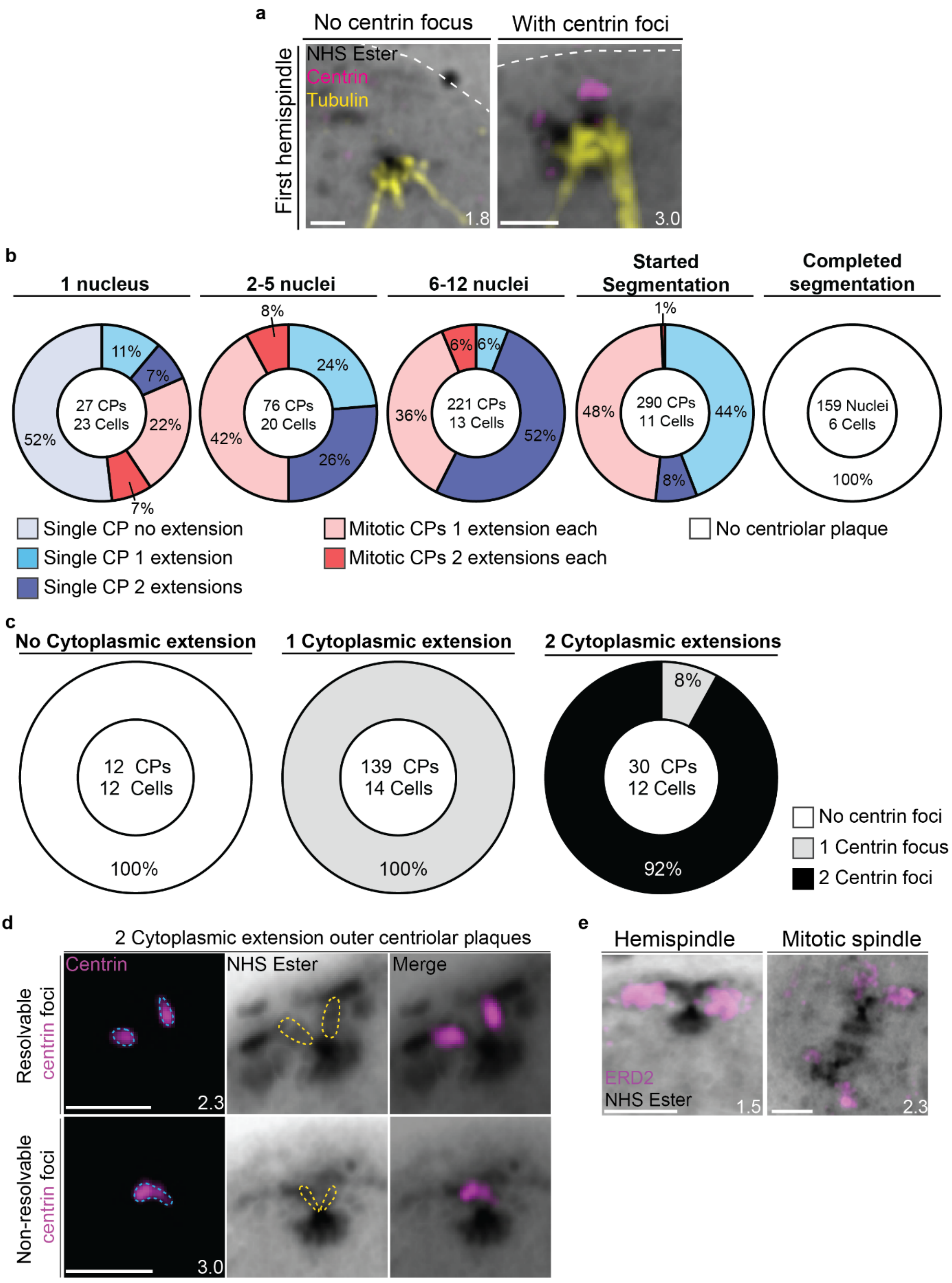


**Figure 2 – Figure Supplement 1**

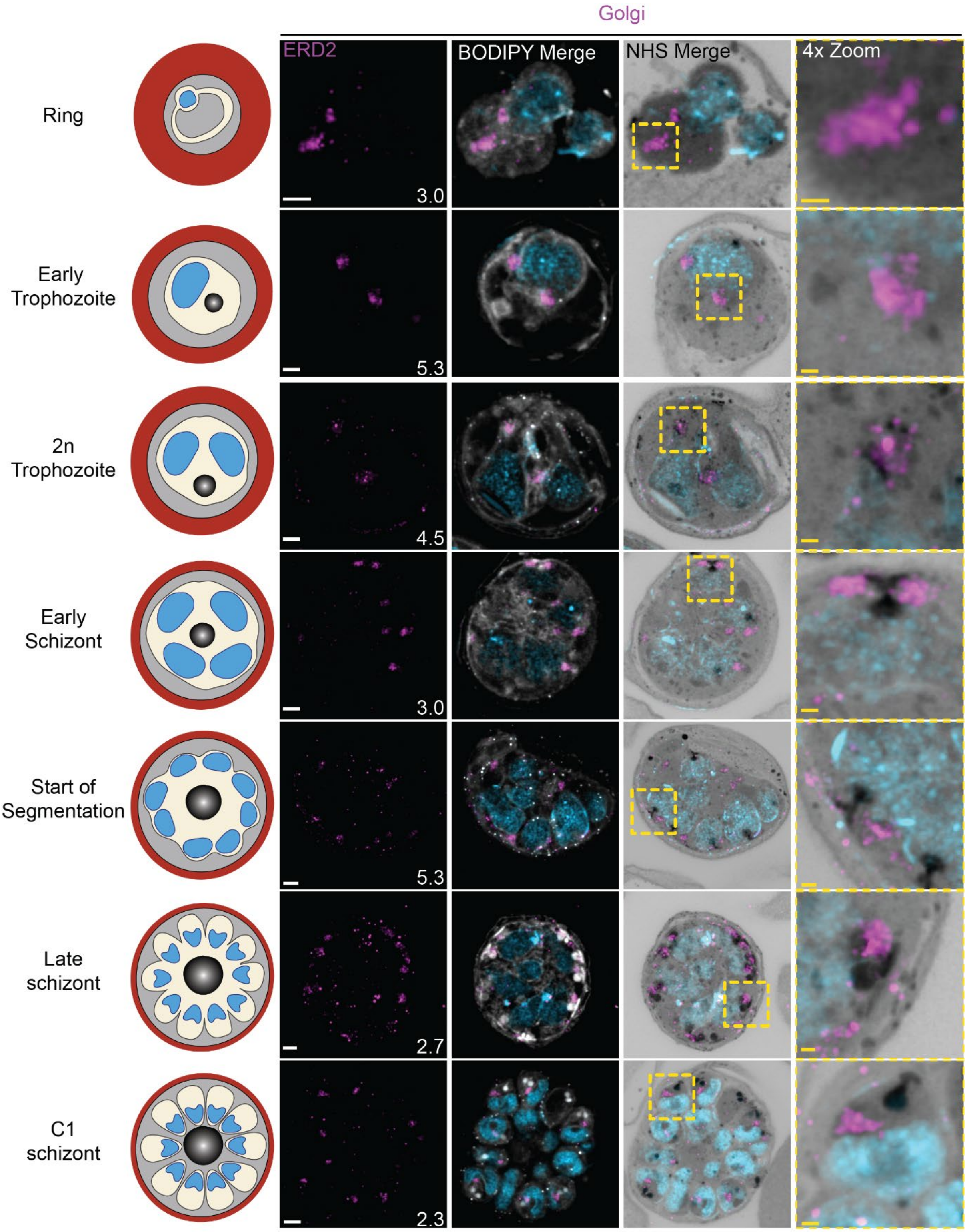


**Figure 2 – Figure Supplement 2**

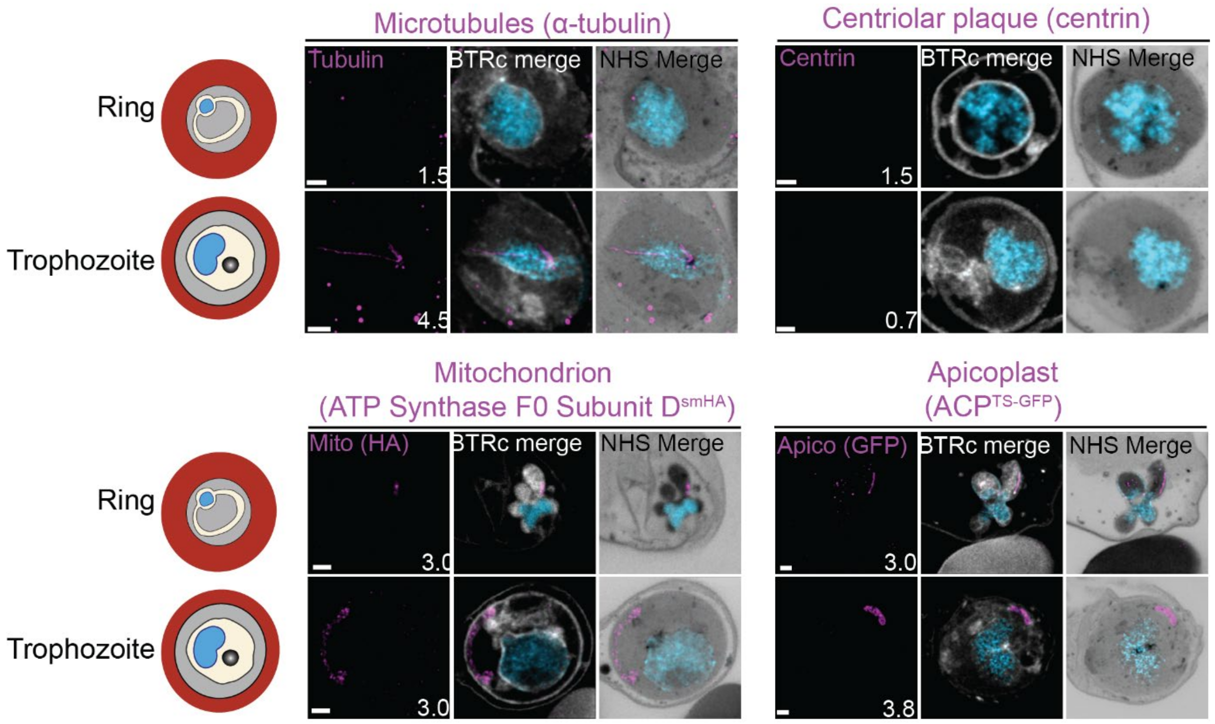


**Figure 3 – Figure Supplement 1**

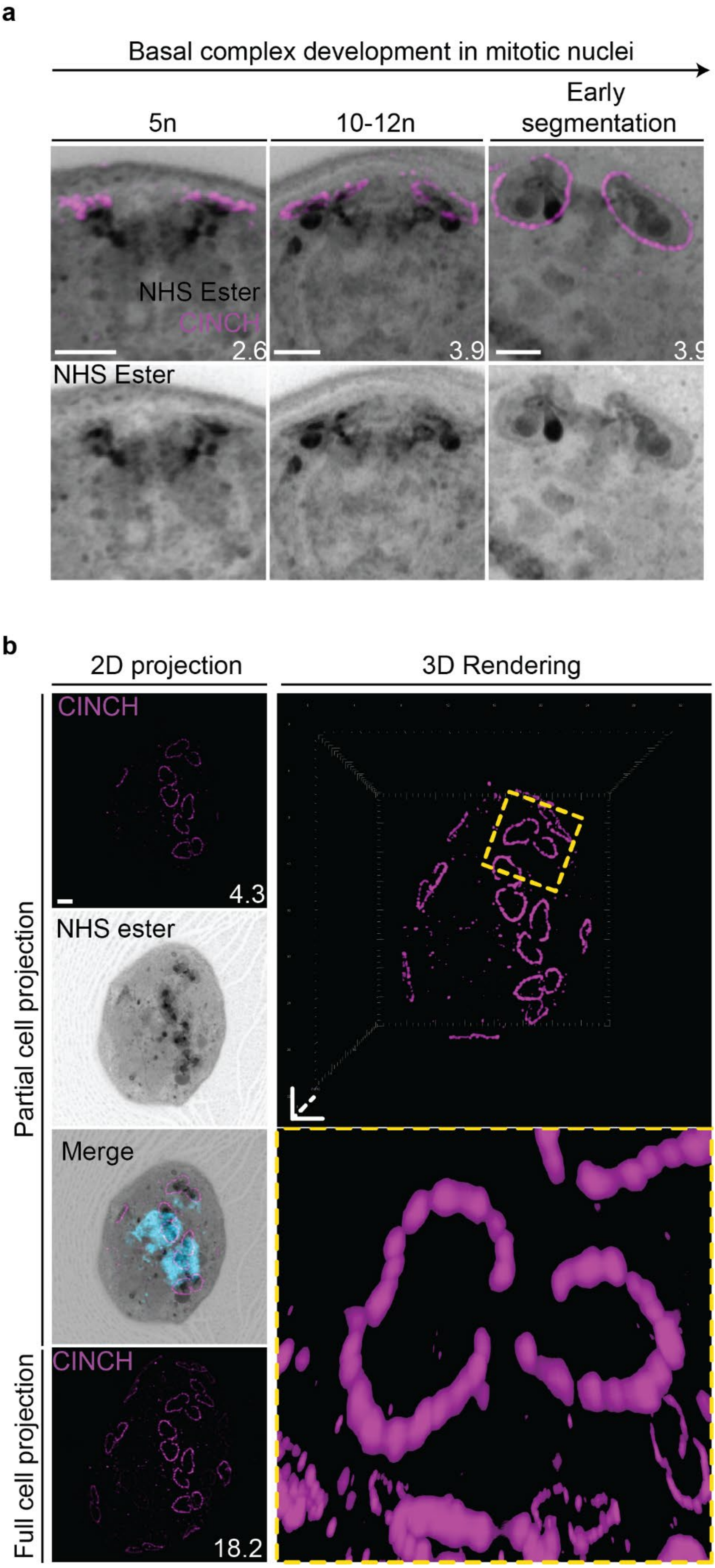


**Figure 4 – Figure Supplement 1**

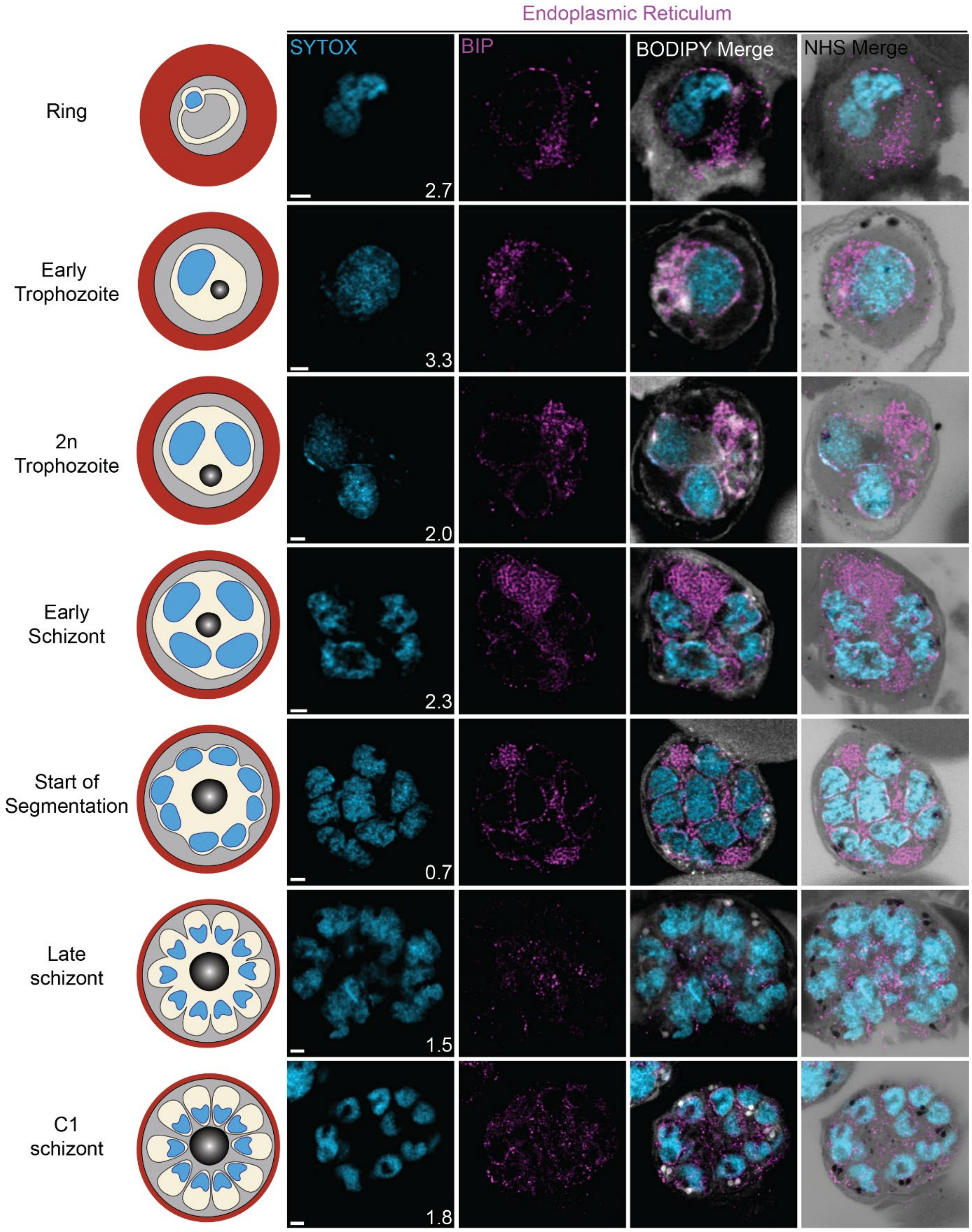


**Figure 4 – Figure Supplement 2**

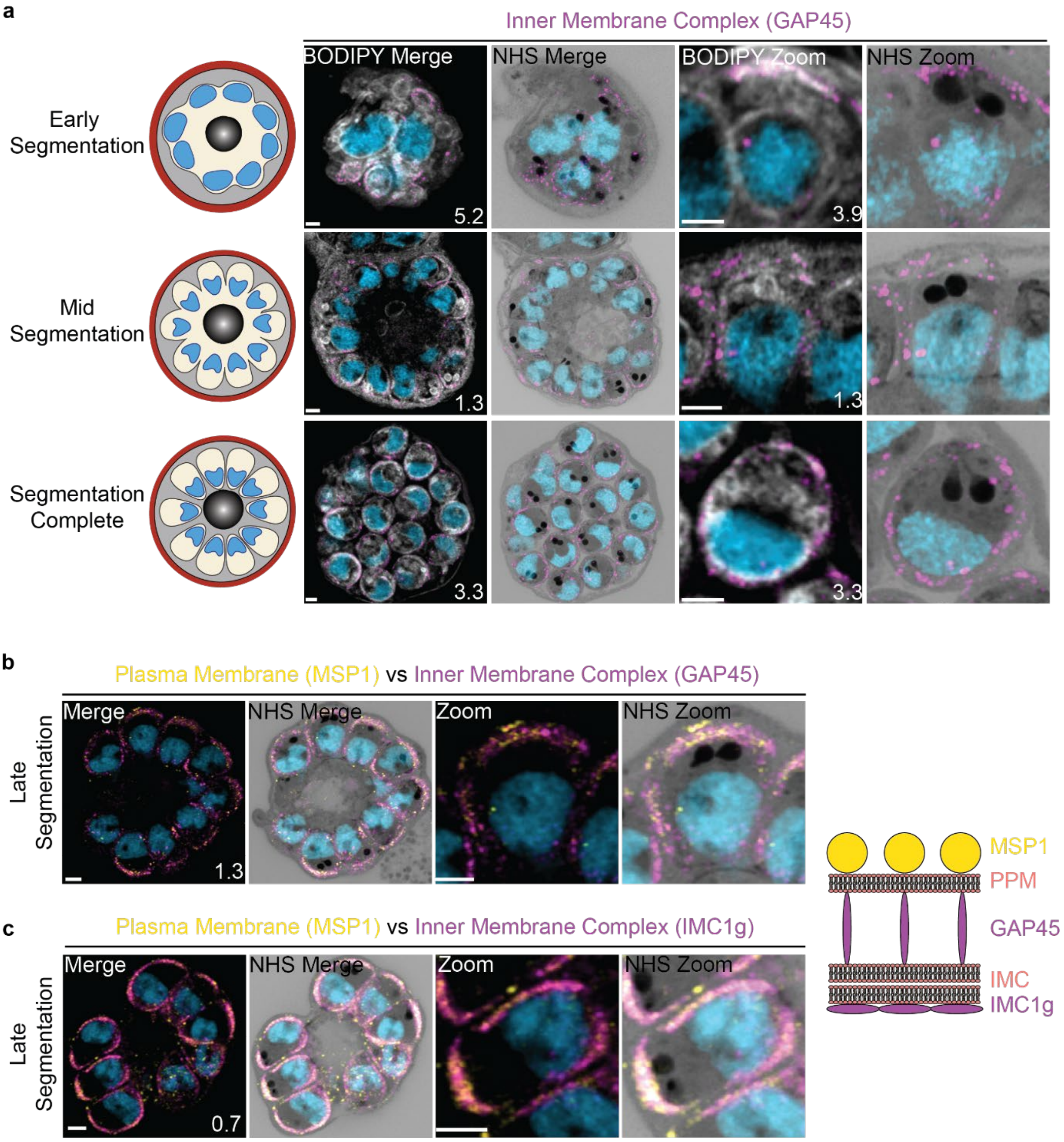


**Figure 4 – Figure Supplement 3**

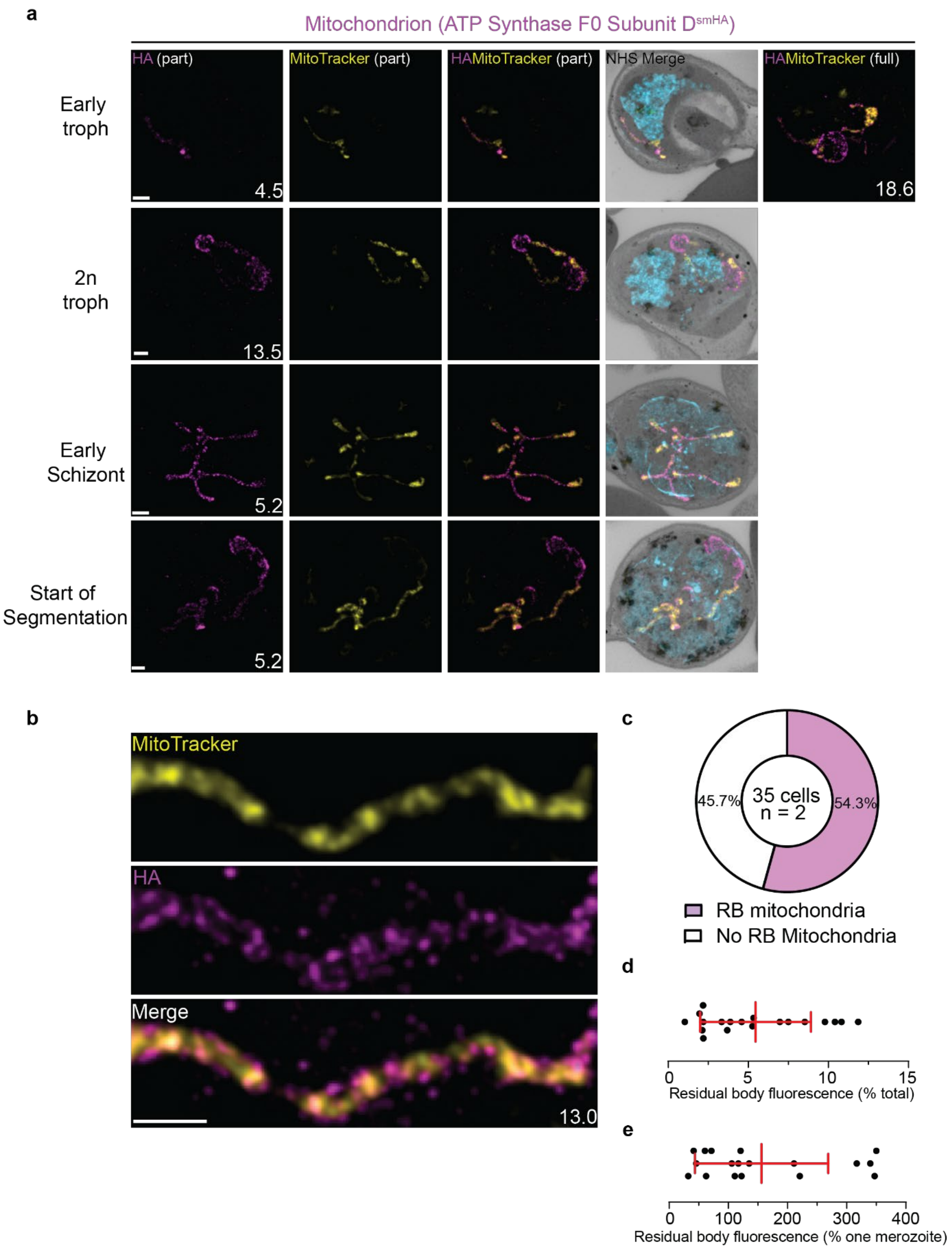


**Figure 5 – Figure Supplement 1**

a

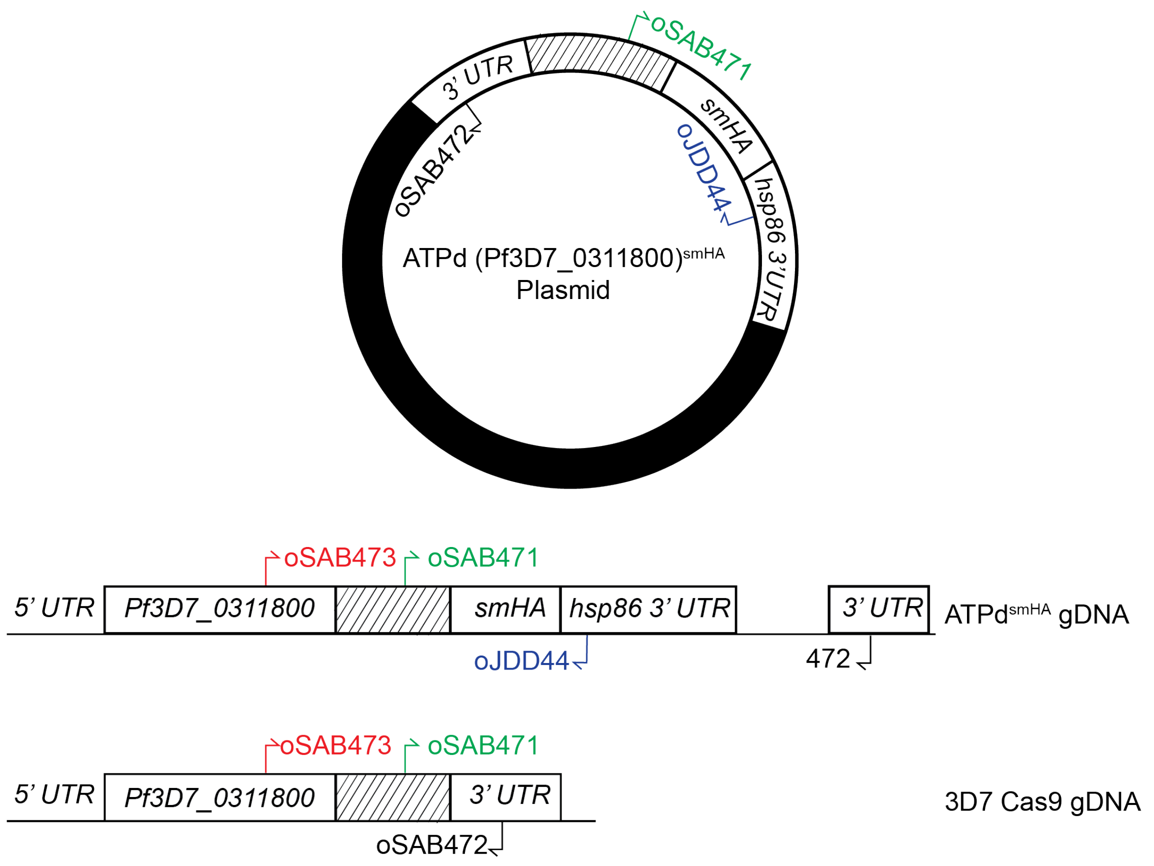


b

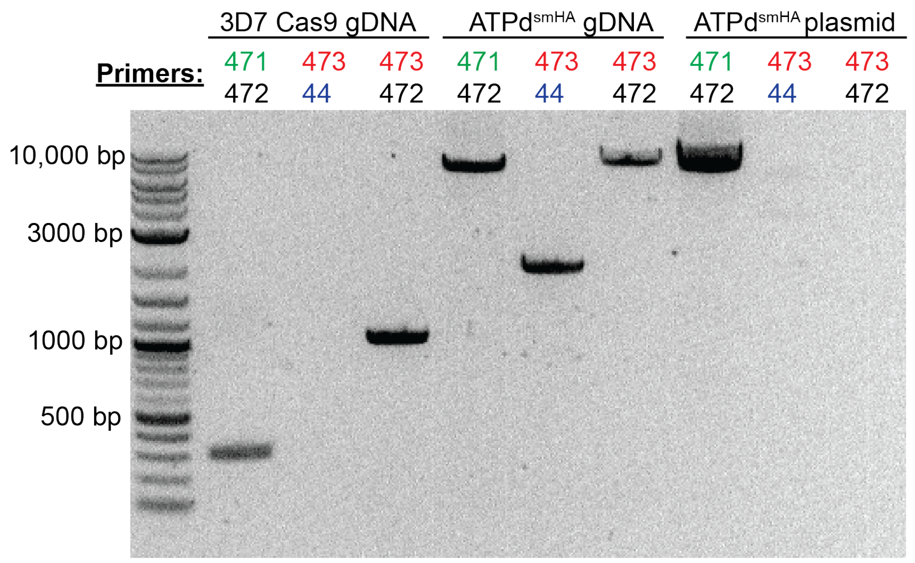


| | 471/472 | 473/44 | 473/472 |
|---|---|---|---|
| 3D7 Cas9 gDNA | 321 bp | -------- | 1038 bp |
| ATPd$^{smHA}$ gDNA | 6890 bp | 2017 bp | 8008 bp |
| ATPd$^{smHA}$ plasmid | 6890 bp | -------- | -------- |

**Figure 5 – Figure Supplement 2**

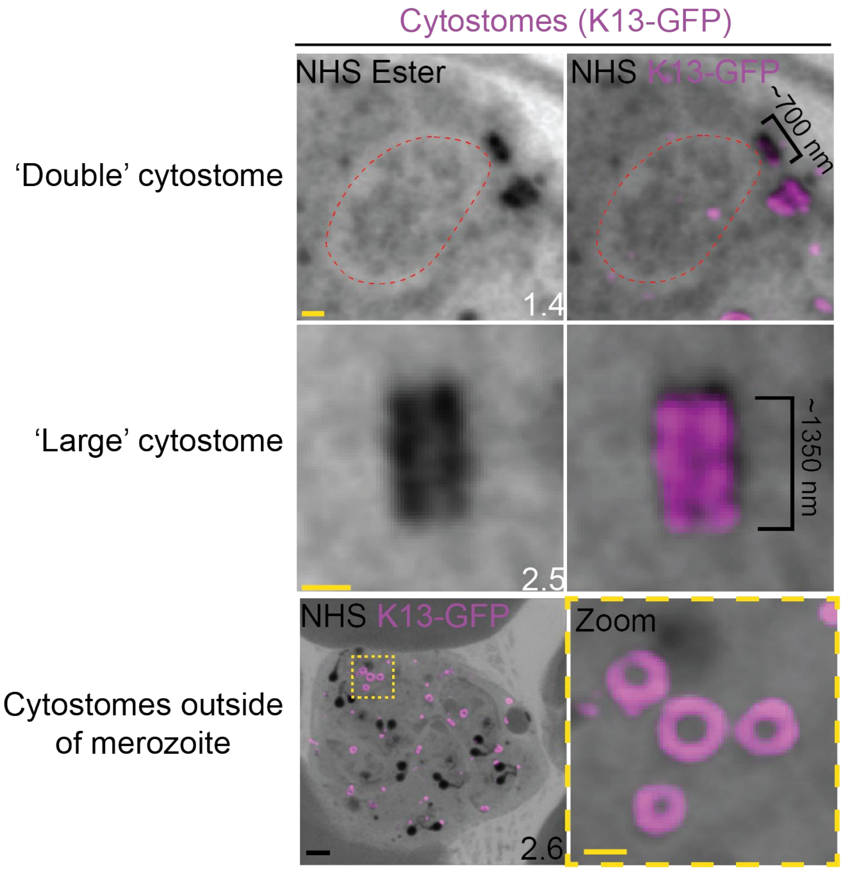


**Figure 7 – Figure Supplement 1**

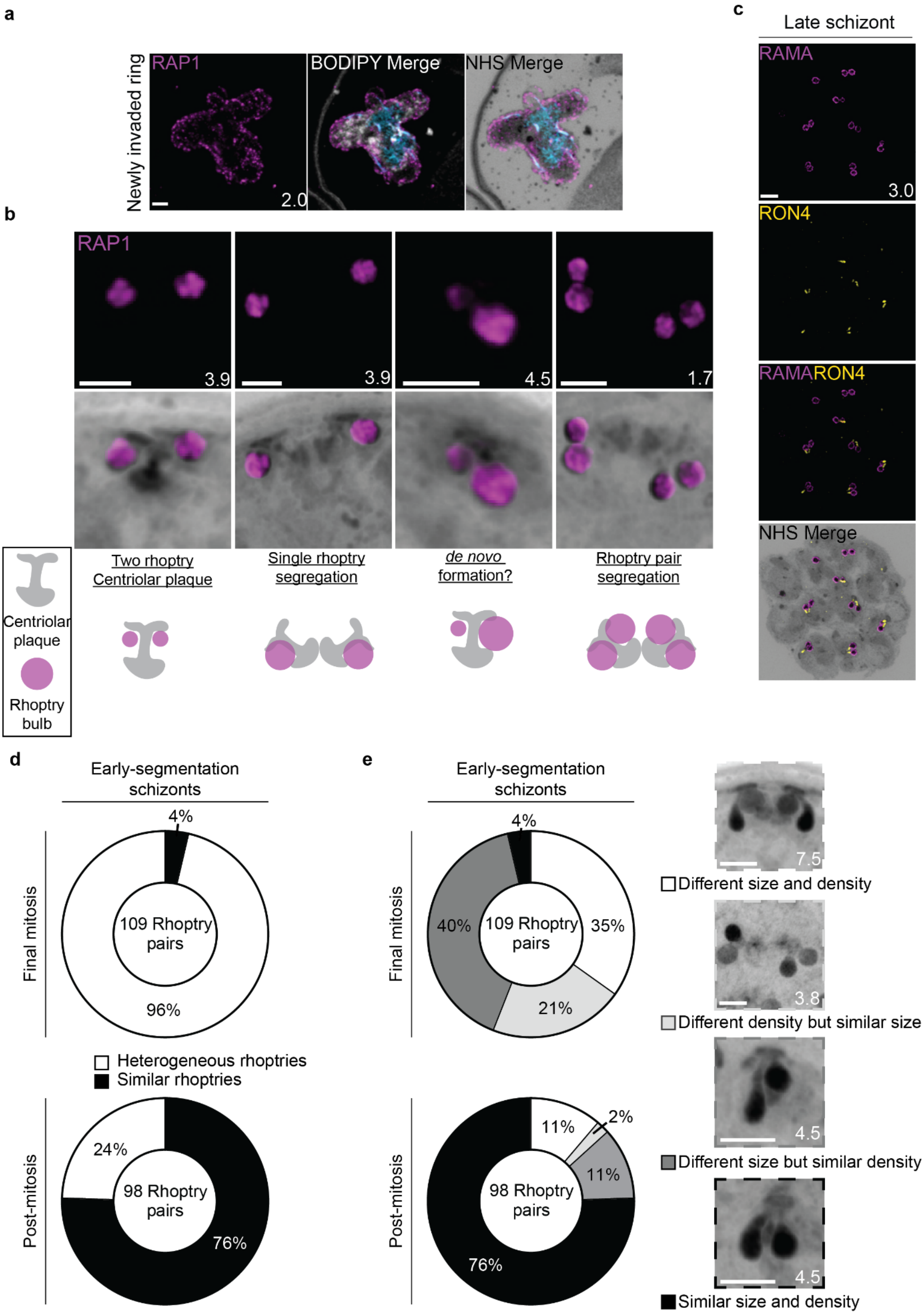


**Figure 8 – Figure Supplement 1**
